# Persons post-stroke restore step length symmetry by walking asymmetrically

**DOI:** 10.1101/799775

**Authors:** Purnima Padmanabhan, Keerthana Sreekanth Rao, Shivam Gulhar, Kendra M. Cherry-Allen, Kristan A. Leech, Ryan T. Roemmich

**Affiliations:** Center for Movement Studies, Kennedy Krieger Institute, Baltimore, MD 21205; Dept of Neuroscience, Johns Hopkins University School of Medicine, Baltimore, MD 21205; Dept of Biomedical Engineering, Johns Hopkins University, Baltimore, MD 21218; Virginia Commonwealth University School of Medicine, Richmond, VA 23298; Dept of Physical Medicine and Rehabilitation, Johns Hopkins University School of Medicine, Baltimore, MD 21205; Division of Biokinesiology and Physical Therapy, University of Southern California, Los Angeles, CA 90007

## Abstract

**Background:** Restoration of step length symmetry is a common rehabilitation goal after stroke. Persons post-stroke often retain the capacity to walk with symmetric step lengths (“symmetric steps”); however, the resulting walking pattern remains effortful. Two key questions with direct implications for rehabilitation have emerged: 1) how do persons post-stroke generate symmetric steps, and 2) why do symmetric steps remain so effortful?

**Objective:** To understand how persons post-stroke generate symmetric steps and how the resulting gait pattern relates to the metabolic cost of transport.

**Methods:** Ten persons post-stroke walked on an instrumented treadmill under two conditions: preferred walking and symmetric stepping (using visual feedback). We recorded kinematic, kinetic, and metabolic data during both conditions.

**Results:** Persons post-stroke restored step length symmetry using energetically expensive, asymmetric patterns. Impaired paretic propulsion and abnormal vertical movement of the center of mass were evident during both preferred walking and symmetric stepping. These deficits contributed to diminished positive work performed by the paretic limb on the center of mass in both conditions. Decreased positive paretic work correlated with increased metabolic cost of transport, decreased self-selected walking speed, and increased asymmetry in limb kinematics.

**Conclusions:** It is important to consider the mechanics used to restore symmetric steps when designing interventions to improve walking after stroke. Facilitating symmetric steps via increased paretic propulsion or enabling paretic limb advancement without excessive vertical movement may enable persons post-stroke to walk with a less effortful, more symmetric gait pattern.

## INTRODUCTION

Gait dysfunction is common after stroke^1^. Persons post-stroke exhibit slow walking speeds^2–4^, gait asymmetry^4,5^, and an elevated metabolic cost of transport (i.e., energy expended per meter walked)^6–8^. Gait training is a key component of stroke rehabilitation, as persons post-stroke frequently list gait improvement among their most desired rehabilitation goals^9^.

Many rehabilitation approaches aim to restore step length symmetry^10–16^. The rationale for restoring step length symmetry is multifaceted: 1) asymmetric stepping increases the cost of transport in healthy adults^17^, 2) persons post-stroke who walk with more asymmetric step lengths also tend to exhibit poorer balance^18^ and more effortful gait patterns^19^, 3) step length is easy to measure and manipulate in clinical settings (e.g.,n “step to the lines on the floor”), and 4) step length asymmetry is a simple, discrete metric that manifests from complex kinematic and kinetic asymmetries that can be difficult to treat in isolation. Consequently, there has been increasing interest in restoring step length symmetry after stroke, especially after recent intervention studies showed that improved step length symmetry coincided with improvements in gait speed^15^ and cost of transport^19^.

However, it is not clear that restoration of step length symmetry alone should lead to improvements in gait speed or cost of transport. Persons post-stroke retain the capacity to walk with improved step length symmetry, even within a single testing session^16,20,21^. But unlike the intervention studies mentioned above, single-session studies have shown cost of transport to be similar whether persons post-stroke walk with asymmetric or symmetric step lengths^16,21^. These findings suggest that improvements in gait speed and cost of transport likely arise from changes in kinematic or kinetic parameters that more directly influence gait speed or energetics and also affect step length symmetry. From this perspective, interventions that aim to restore step length symmetry but do not affect these critical underlying factors may not result in meaningful gait improvement. The ability to lessen cost of transport with an intervention aiming to restore step length symmetry likely depends on 1) the underlying causes of the asymmetry (which vary from person to person^21,22^), and 2) the gait mechanics used to generate the symmetric step lengths.

Here, we aimed to understand how persons post-stroke changed their walking patterns to restore step length symmetry and how these gait mechanics related to the cost of transport. We asked: do persons post-stroke restore step length symmetry by restoring symmetric gait mechanics or by relying on asymmetric compensatory mechanics? We hypothesized that persons post-stroke would restore step length symmetry using asymmetric walking patterns. We then aimed to explain why these asymmetric gait patterns cost so much energy despite restoration of step length symmetry.

## METHODS

### General methods

Ten persons post-stroke participated (7M/3F, age (mean ±SEM): 56±4 years, lower extremity Fugl-Meyer^23^: 26±1, body mass: 91±6 kg, all >6 months post-stroke). Participants reported no additional neurological, musculoskeletal, or cardiovascular conditions. We determined preferred walking speed as the average speed of three overground ten-meter walk tests (0.81±0.09 m/s, range: 0.40-1.25 m/s). Eight participants held onto the treadmill handrails, two wore ankle-foot orthoses, and one received functional electrical stimulation of the tibialis anterior. All participants wore a safety harness that did not provide body weight support, provided written informed consent in accordance with the Johns Hopkins Medicine Institutional Review board prior to participation, and received monetary compensation.

We recorded kinematic (100 Hz) and kinetic (1000 Hz) data using a three-dimensional motion capture system (Vicon, Oxford, UK) and instrumented split-belt treadmill (Motek, Amsterdam, NL; Figure 1A). We placed retroreflective markers over the seventh cervical vertebrae, tenth thoracic vertebrae, jugular notch, xiphoid process, and bilaterally over the second and fifth metatarsal heads, calcaneus, medial and lateral malleoli, shank, medial and lateral femoral epicondyles, thigh, greater trochanter, iliac crest, and anterior and posterior superior iliac spines. We filtered marker trajectory and ground reaction force (GRF) data with fourth order low-pass Butterworth filters (6 Hz and 15 Hz cut-off frequencies, respectively). GRF magnitudes were set to zero if the vertical GRF magnitude was less than 32 N. All participants wore comfortable shoes and form-fitting clothing.

**Figure 1.**
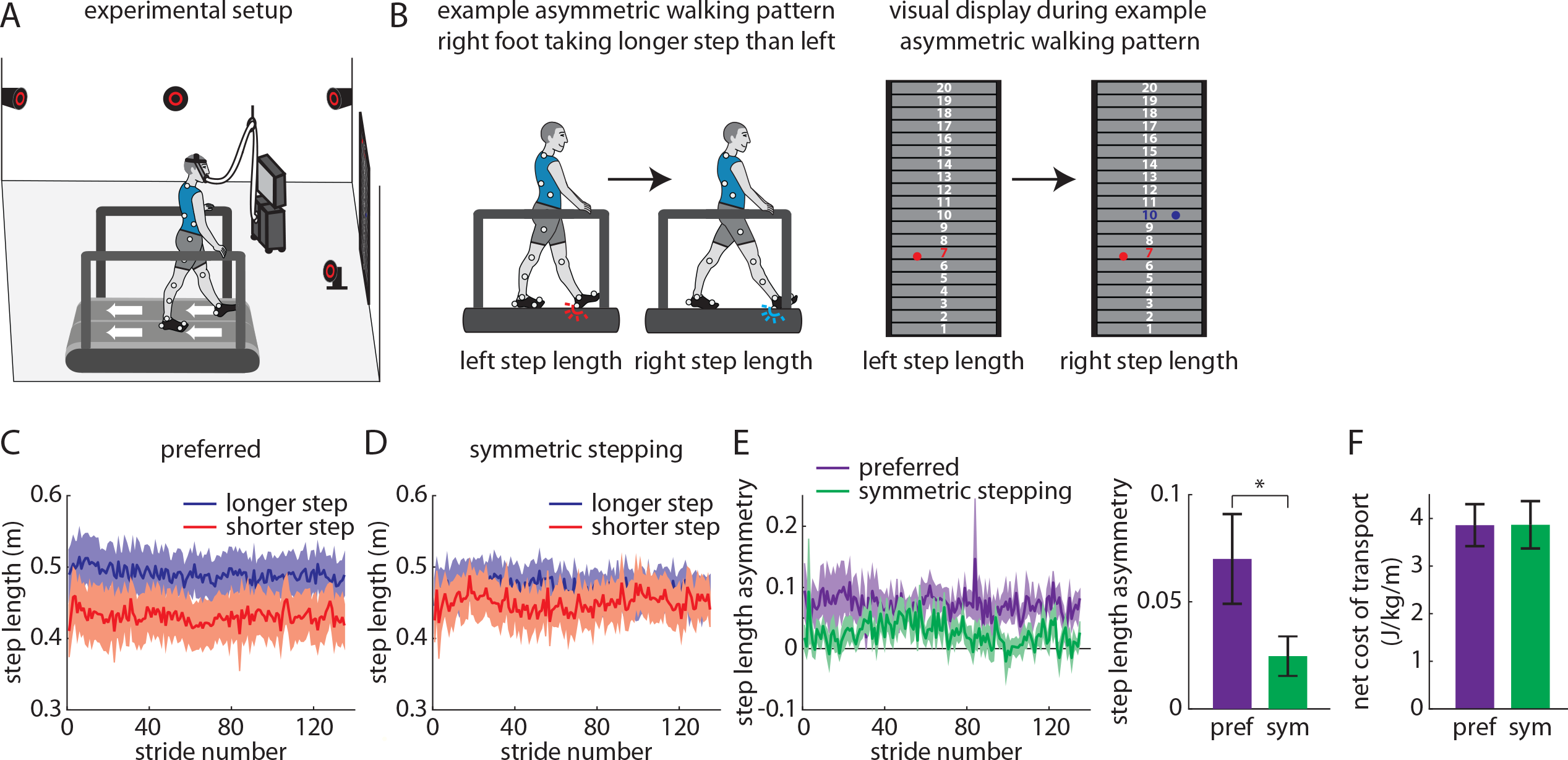
*A*) Experimental setup. *B*) Example participant walking with asymmetric step lengths (right longer than left) and resulting visual display showing step length feedback bilaterally. *C* and *D*) Step lengths for the limbs that took longer (blue) and shorter (red) steps at baseline during *C*) preferred walking and *D*) symmetric stepping. *E*) Step length asymmetry decreases significantly during symmetric stepping (green) as compared to preferred walking (purple). *F*) The net metabolic cost of transport is similar between preferred walking and symmetric stepping.

We collected metabolic data using a TrueOne 2400 system (Parvomedics, Sandy, UT) that warmed up for >30 minutes before data collection and was calibrated to manufacturer specifications. We sampled oxygen consumption and carbon dioxide production breath-by-breath. We collected two minutes of baseline metabolic data during quiet standing. We used a traditional equation^24^ to calculate total metabolic power during walking trials and subtracted baseline metabolic power to calculate net metabolic power. We calculated net metabolic cost of transport (herein referred to as cost of transport) by normalizing net metabolic power to treadmill speed.

### Visual display

Our feedback display showed 20 vertically-arranged virtual targets on a 4 m × 2.5 m screen in front of the treadmill^16,25^. The virtual targets provided a reference frame for participant step lengths. Red and blue circles appeared on the left and right halves of the display, respectively, at heel-strikes (detected in real-time using the force plates under each treadmill belt) to represent left and right step lengths (Figure 1B). The white number centered inside the target changed color (red at left heel-strike, blue at right heel-strike) when it had been reached with each foot.

### Protocol

Participants performed three treadmill walking trials at preferred speeds. Participants first walked for four minutes without feedback (baseline). This enabled us to measure baseline asymmetry and identify which leg took a longer step. We then displayed step length feedback, and participants walked with either their preferred gait pattern or symmetric step lengths (order randomized). During preferred walking, we asked participants to walk normally while receiving visual feedback. During symmetric stepping, we asked participants to hit the same target with each pair of steps. We did not enforce constraints on individual step lengths or provide instructions about which leg should step longer or shorter to restore step length symmetry.

### Spatiotemporal and kinematic measurements

We measured step length as the distance between the lateral malleoli markers along the anterior-posterior axis at heel-strike and step length asymmetry as the difference in consecutive step lengths between the leg that took a longer step at baseline and the leg that took shorter step normalized to their sum:

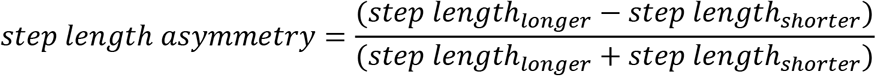

We also developed a measure of kinematic asymmetry (“interlimb asymmetry”) that was agnostic to each participant’s idiosyncratic deficits. This was important in enabling us to understand whether the participants restored symmetric step lengths with asymmetric kinematics without assigning the asymmetries to specific joints. Furthermore, we could determine whether the kinematic patterns used to restore symmetric step lengths were similar to the participants’ preferred walking patterns.

Interlimb asymmetry quantifies asymmetry in individual limb segment contributions to step length. Consider a right step length of 0.5 m. If the distance between the left lateral malleolus and lateral femoral epicondyle markers is 0.05 m at this right heel-strike, the trailing (left) shank segment contribution to right step length is 0.05 m/0.5 m, or 0.10. These segment contributions were calculated along the anterior-posterior axis for the following segments and sum to 1:

- trailing shank (trailing lateral malleolus to lateral femoral epicondyle, *a*)
- trailing thigh (trailing lateral femoral epicondyle to greater trochanter, *b*)
- trailing pelvis (trailing greater trochanter to iliac crest, *c*)
- trailing contribution from pelvic rotation (trailing iliac crest to center of pelvis, *d*)
- leading contribution from pelvic rotation (center of pelvis to leading iliac crest, *e*)
- leading pelvis (leading iliac crest to greater trochanter, *f*)
- leading thigh (leading greater trochanter to lateral femoral epicondyle, *g*)
- leading shank (leading lateral femoral epicondyle to lateral malleolus, *h*)

We calculated interlimb asymmetry by summing the segment asymmetries between the left (*l*) and right (*r*) legs at consecutive heel-strikes (i.e., absolute values of the differences between each segment contribution bilaterally):

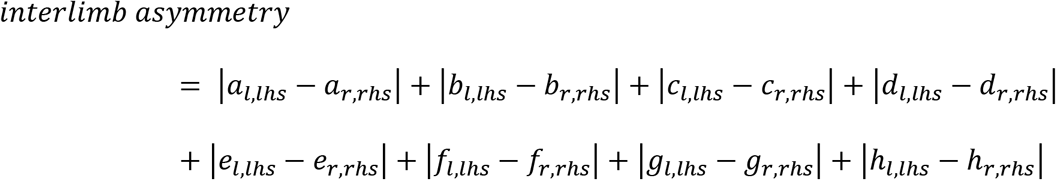

Interlimb asymmetry is bounded between 0 (completely symmetric segment contributions) and 2 (completely asymmetric segment contributions). Importantly, this metric can remain constant across different step lengths as long as the segment contributions scale proportionally.

### Kinetic measurements

We calculated the instantaneous center of mass (COM) mechanical power using the individual limbs method^26^ as has been done previously in persons post-stroke^27,28^. This method assumes a mechanical model of gait that allows for calculation of instantaneous COM power generated by each leg as the dot product of the GRF vector of each leg and the COM velocity vector^26^. We performed these calculations for the last five strides of each condition that exhibited clean force plate strikes (i.e., each foot landed on a different force plate). Each stride began with a paretic limb heel-strike. We partitioned each stride into four periods (two step-to-step transition periods and two non-transition periods) based on time points when the COM velocity vector was redirected within the sagittal plane^29^. The onset of the first period was considered the cessation of the fourth period of the previous stride. We calculated the positive and negative work done on the COM by each limb during the four periods by integrating the positive and negative portions of the power curve within each period. All kinetic measures were normalized to body mass.

### Data analysis

We averaged step length asymmetry, interlimb asymmetry, and segment asymmetries across the four-minute trial for each participant and condition. We calculated cost of transport over the final minute of each trial to ensure steady state measurement. In our GRF and COM velocity analyses, we set the anterior-posterior (AP) axis to be positive in the forward direction, mediolateral (ML) axis to be positive in the direction from the paretic limb toward the nonparetic limb, and vertical axis to be positive upward. We calculated GRF peaks as the most positive values produced along each axis by each leg (except the nonparetic ML GRF peak, which was calculated as the most negative value) stride-by-stride over the final five clean strides. We identified peak COM velocities at two time points. We calculated peak AP and vertical COM velocities as the most positive velocities observed when the corresponding GRF magnitude was also positive. We calculated peak ML COM velocities as the most positive COM velocity when the paretic ML GRF was positive (paretic) and the most negative velocity when the nonparetic ML GRF was negative (nonparetic). We calculated positive and negative work done by each limb during each of the four gait cycle periods stride-by-stride over the final five clean strides and then averaged across strides for each participant and condition. We also calculated total positive and negative work done by each limb across the gait cycle stride-by-stride over the final five clean strides and then averaged across strides for each participant and condition.

### Statistical analysis

We performed paired t-tests to compare step length asymmetry, cost of transport, and interlimb asymmetry between conditions (preferred walking and symmetric stepping). We performed a 7×2 limb segment (*a* through *h* as described above, with *d* and *e* summed to indicate pelvic rotation) × condition repeated measures ANOVA to compare segment asymmetry among segments and between conditions. We performed 2×2 leg (paretic, nonparetic) × condition repeated measures ANOVAs to compare GRF peaks, COM velocity peaks, and positive and negative work done across legs and conditions. We performed Pearson’s correlations to assess the following relationships: interlimb asymmetry during preferred walking vs. interlimb asymmetry during symmetric stepping, interlimb asymmetry vs. cost of transport, cost of transport vs. positive paretic and nonparetic work, preferred walking speed vs. positive paretic and nonparetic work, and interlimb asymmetry vs. positive paretic and nonparetic work. We used an alpha level of 0.05, performed Mauchly’s tests of sphericity (Greenhouse-Geisser corrections were applied when sphericity was violated), and applied post hoc corrections for multiple comparisons where appropriate (Bonferroni for analyses with three comparisons, Dunn-Sidak otherwise).

## RESULTS

### Persons post-stroke can walk with more symmetric step lengths, but this does not change the cost of transport

As expected, participants walked with asymmetric step lengths during preferred walking (Figure 1C) and successfully adjusted their step lengths during the symmetric stepping condition (Figure 1D) to reduce step length asymmetry (t(9)=3.41, p<0.01; Figure 1E). We replicated prior findings^16,21^ showing that restoring step length symmetry with visual feedback had no significant effect on cost of transport (t(9)=−0.05, p=0.96; Figure 1F).

### Persons post-stroke exhibit marked interlimb asymmetry even when walking with symmetric step lengths

A conceptual illustration of how we expected interlimb asymmetry may differ between healthy symmetric walking and symmetric stepping after stroke is shown in Figures 2A and 2B. Briefly, we hypothesized that healthy walking consists of symmetric step lengths and similar contributions of each segment to step lengths bilaterally, resulting in low interlimb asymmetry (Figure 2A). On the contrary, we expected that symmetric stepping after stroke consists of symmetric step lengths but asymmetric segment contributions, resulting in high interlimb asymmetry (Figure 2B).

**Figure 2.**
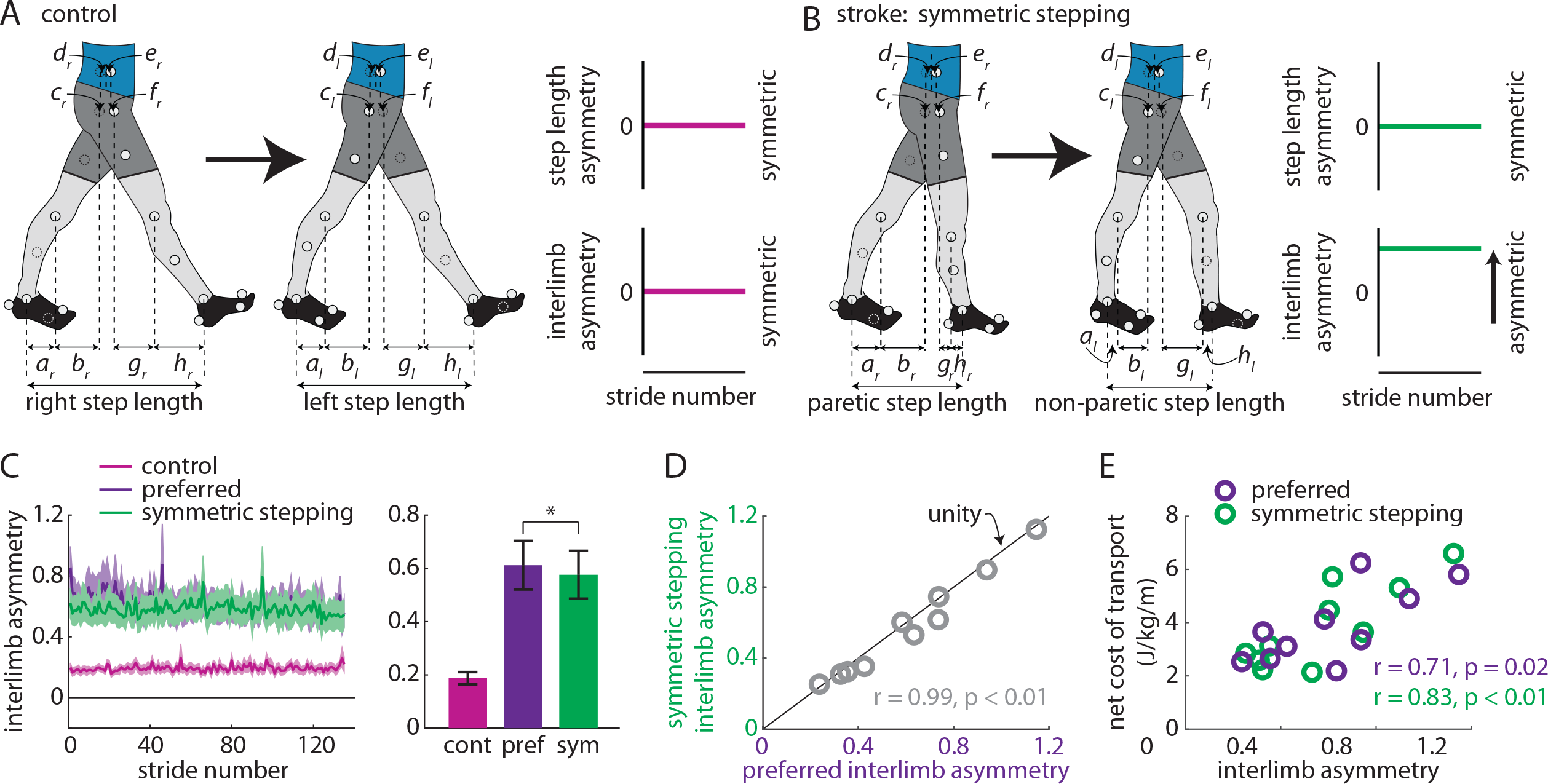
*A* and *B*) Conceptual illustrations of hypotheses regarding walking patterns used by control (healthy young adults) and persons post-stroke to achieve step length symmetry. *A*) Healthy adults achieve step length symmetry using symmetric kinematics (as represented by small interlimb asymmetry). *B*) Persons post-stroke achieve step length symmetry using asymmetric kinematics (large interlimb asymmetry). *C*) Persons post-stroke walk with marked interlimb asymmetry during preferred walking and symmetric stepping (mean ± SEM; * p<0.05 between conditions). Control (healthy young adult) data shown for reference (not included in statistical analyses). *D*) Interlimb asymmetry during symmetric stepping correlates strongly with interlimb asymmetry during preferred walking in persons post-stroke. *E*) The net metabolic cost of transport correlates strongly with interlimb asymmetry during preferred walking and symmetric stepping.

We observed a significant reduction in interlimb asymmetry during symmetric stepping as compared to preferred walking (t(9)=2.34, p=0.04; Figure 2C). While this comparison reached statistical significance, interlimb asymmetry during symmetric stepping remained markedly increased when compared to healthy gait (for reference, data from eight healthy adults (age: 26±5 years) walking at 1.25 m/s shown in Figure 2C). Interlimb asymmetry during preferred walking correlated strongly with interlimb asymmetry during symmetric stepping (r=0.99, n=10, p<0.01; Figure 2D) and, qualitatively, the data fell near the unity line (Figure 2D), suggesting that persons post-stroke showed similar interlimb asymmetry during preferred walking and symmetric stepping. Interlimb asymmetry was significantly associated with cost of transport during preferred walking (r=0.71, n=10, p=0.02) and symmetric stepping (r=0.83, n=10, p<0.01; Figure 2E), revealing that kinematic asymmetries are related to cost of transport regardless of step length asymmetry.

We next considered that interlimb asymmetry could remain similar across conditions while individual segment asymmetries could be reorganized. We did not find this to be the case. We show the limb segment orientations during representative steps of symmetric stepping for each participant (Figure 3A). We compared the individual segment asymmetries (e.g., |*a_l,lhs_* − *a_r,rhs_*|) across segments and between conditions. ANOVA revealed a significant main effect of segment (F(1,9)=4.84, p=0.013). Post hoc analyses revealed that segment asymmetry was significantly larger in pelvic rotation (*d+e*) than trailing pelvis translation (*c*) (p=0.036). We did not observe a significant main effect of condition (F(1,9)=2.69, p=0.135; Figure 3B) or segment × condition interaction (F(1,9)=0.78, p=0.46). Figures 3C and 3D show how the segment asymmetries contribute to interlimb asymmetry for each participant during each condition. When we compared segment asymmetries after ordering them by which contributed most-to-least strongly to interlimb asymmetry (during preferred walking) between conditions, we also did not observe a significant main effect of condition (F(1,9)=2.69, p=0.135; Figure 3E) or segment × condition interaction (F(1,9)=1.93, p=0.175). As expected, we observed a significant main effect of segment (F(1,9)=23.9, p<0.001).

**Figure 3.**
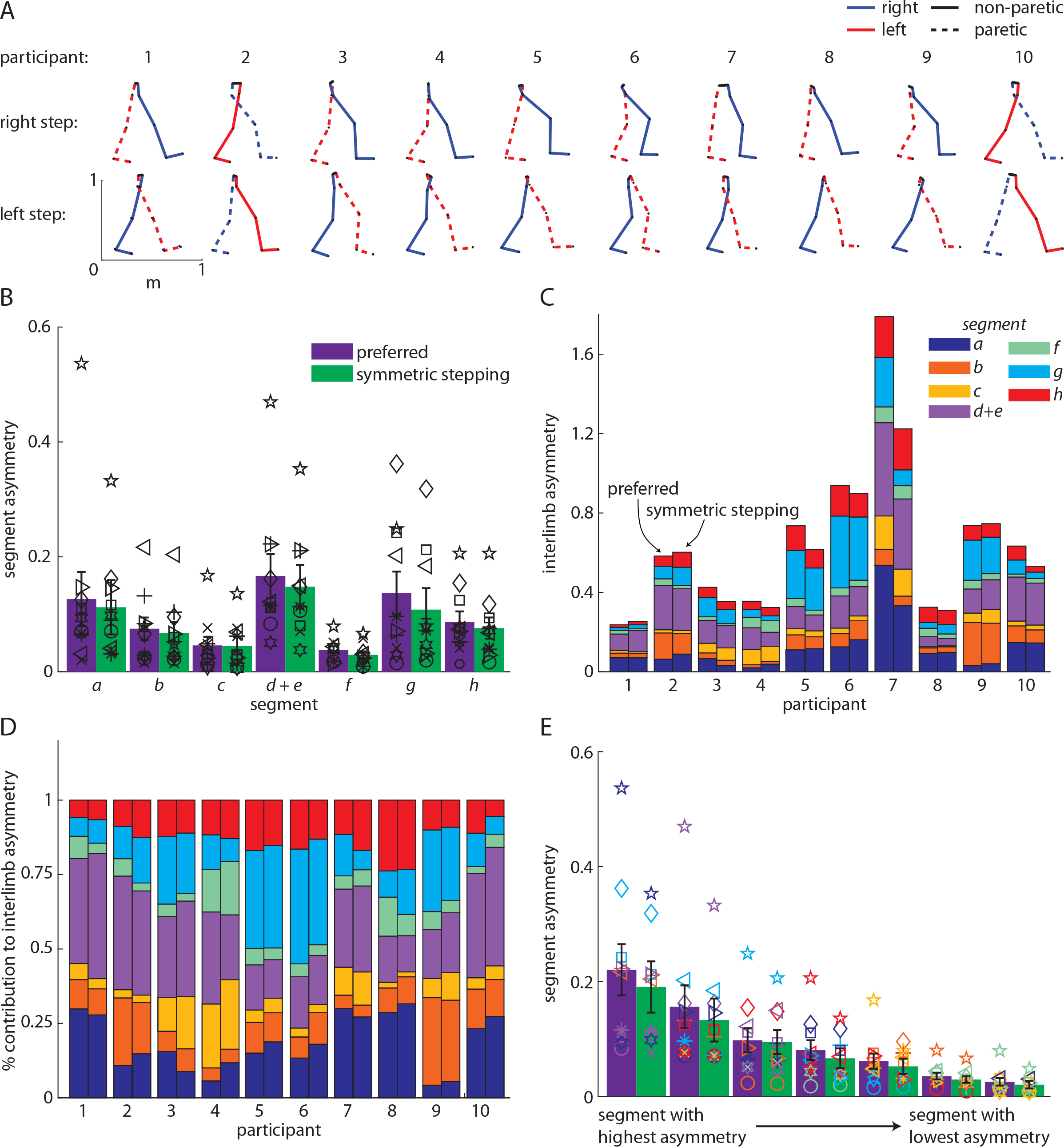
*A*) Limb orientations (blue = right, red = left, solid = non-paretic, dashed = paretic) during representative steps for each participant. *B*) Segment asymmetries during preferred walking and symmetric stepping (organized by segment; mean ± SEM). Symbols represent individual participants. *C*) Segment asymmetry contributions to interlimb asymmetry during preferred walking and symmetric stepping (organized by participant). *D*) Segment asymmetry contributions to interlimb asymmetry during preferred walking and symmetric stepping shown as a percentage of interlimb asymmetry. *E*) Segment asymmetry contributions to interlimb asymmetry during preferred walking and symmetric stepping (segments organized from highest asymmetry to lowest asymmetry). Symbols follow same scheme as in *B* and colors follow the same scheme as in *C* and *D*.

### Asymmetries in AP GRFs, ML GRFs, and vertical COM velocities observed during preferred walking persist during symmetric stepping

We then aimed to understand the features of these asymmetric walking patterns that influence the elevated cost of transport regardless of step length asymmetry. We investigated whether these features were similar in both preferred walking and symmetric stepping, or whether the costs of transport were similarly high in these conditions but resulted from different underlying mechanics. Asymmetric kinematics at heel-strike should result in asymmetric mechanical work done on the COM by each leg, and previous studies demonstrated that mechanical work done on the COM is related to cost of transport in healthy adults^26,30^. Furthermore, prior studies have identified periods of the gait cycle where excessive positive work is often observed post-stroke, contributing to an elevated mechanical energetic cost^7,8,31^.

We investigated the GRF and COM velocity profiles between legs and conditions, as these contribute to the work done over the gait cycle (Figures 4A and 4B). ANOVA revealed a main effect of leg on the AP GRF peak (F(1,9)=10.97, p<0.01), ML GRF peak (F(1,9)=9.71, p=0.01), and vertical COM velocity peak (F(1,9)=7.68, p=0.02). Post hoc analyses revealed that the AP GRF peak was significantly larger in the nonparetic leg than the paretic leg (p<0.01), the ML GRF peak was significantly larger in the paretic leg than the nonparetic leg (p=0.01), and the vertical COM velocity peak was significantly larger during paretic late stance as compared to nonparetic late stance (p=0.02). There were no significant effects of leg on the vertical GRF peak (F(1,9)=1.25, p=0.29), AP COM velocity peak (F(1,9)=4.29, p=0.07), ML COM velocity peak (F(1,9)=2.90, p=0.12). We did not observe significant effects of condition on GRF or COM velocity variables (all p>0.08) or leg × condition interactions (all p>0.41).

**Figure 4.**
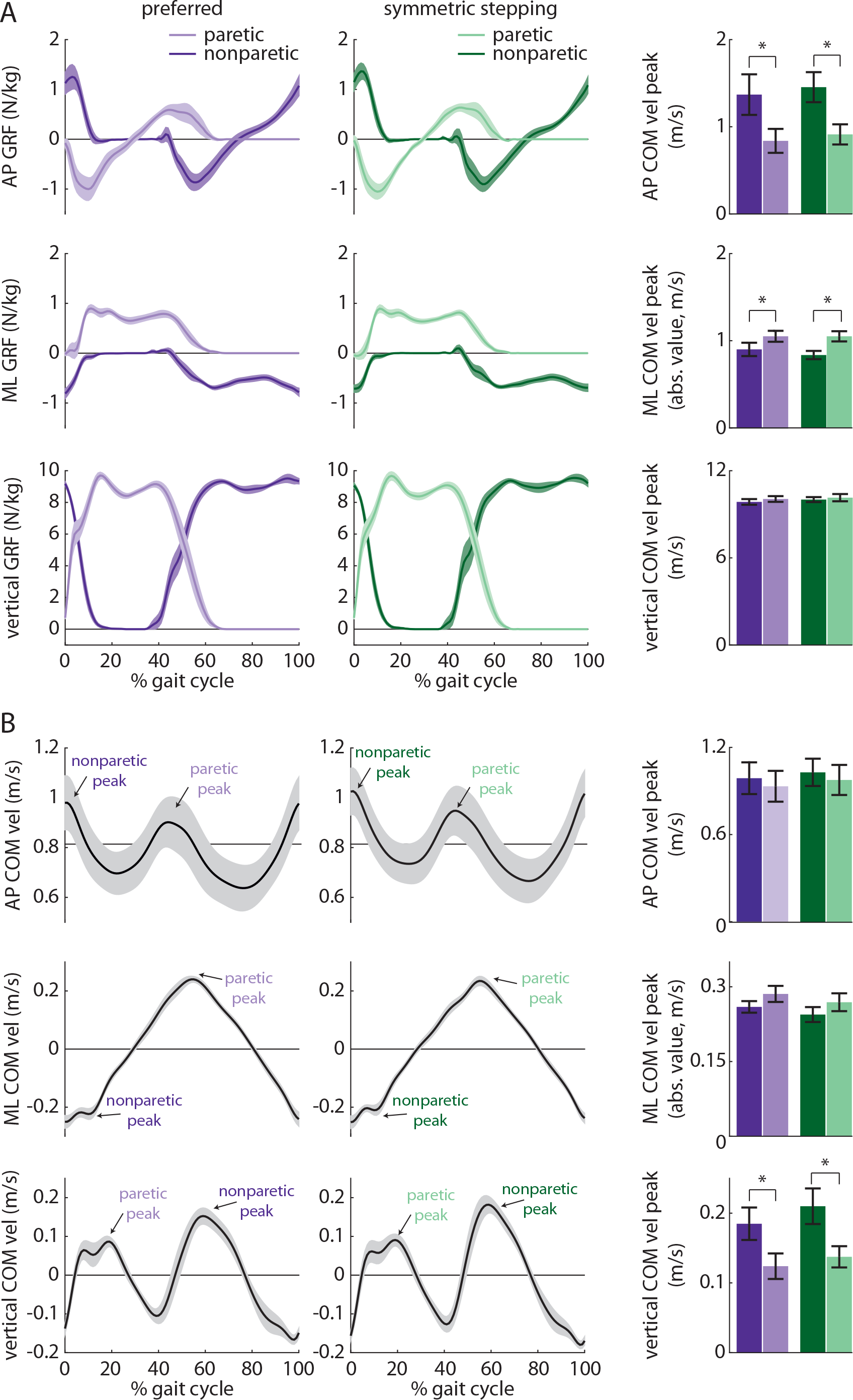
*A*) Anterior-posterior (AP; top), mediolateral (ML; middle), and vertical (bottom) ground reaction force (GRF) profiles for the paretic (light colors) and nonparetic (dark colors) limbs during preferred walking (purple) and symmetric stepping (green). The gait cycle is aligned to paretic heel-strike. Persons post-stroke show decreased peak AP force production and increased peak ML force production in the paretic limb during both conditions. *B*) AP (top), ML (middle), and vertical (bottom) center of mass (COM) velocity profiles during preferred walking and symmetric stepping. Persons post-stroke show increased vertical COM velocity during late paretic stance as compared to late nonparetic stance during both conditions. *A* and *B* show mean ± SEM. * p<0.05 between limbs. We did not observe any significant differences between conditions.

### The nonparetic leg does more positive work than the paretic leg during preferred walking and symmetric stepping

We next investigated the work done on the COM by each leg across conditions. We first calculated COM power for each leg during preferred walking and symmetric stepping (Figures 5A and 5B). We calculated COM work by integrating COM power over each of the four time periods described in the methods (Figures 5C-5F). ANOVA revealed a significant main effect of leg on positive work done 1) by the paretic leg during the first period (step-to-step transition, nonparetic leg trailing) vs. the nonparetic leg during the third period (step-to-step transition, paretic leg trailing; F(1,9)=10.46, p=0.01), and 2) by the paretic leg during the second period (paretic single support) vs. the nonparetic leg during the fourth period (nonparetic single support; F(1,9)=14.10, p<0.01). Post hoc analyses revealed that the nonparetic leg did significantly more positive work during the third period than the paretic leg did during the first period (p=0.01). The nonparetic leg also did significantly more positive work during the fourth period than the paretic leg did during the second period (p<0.01).

**Figure 5.**
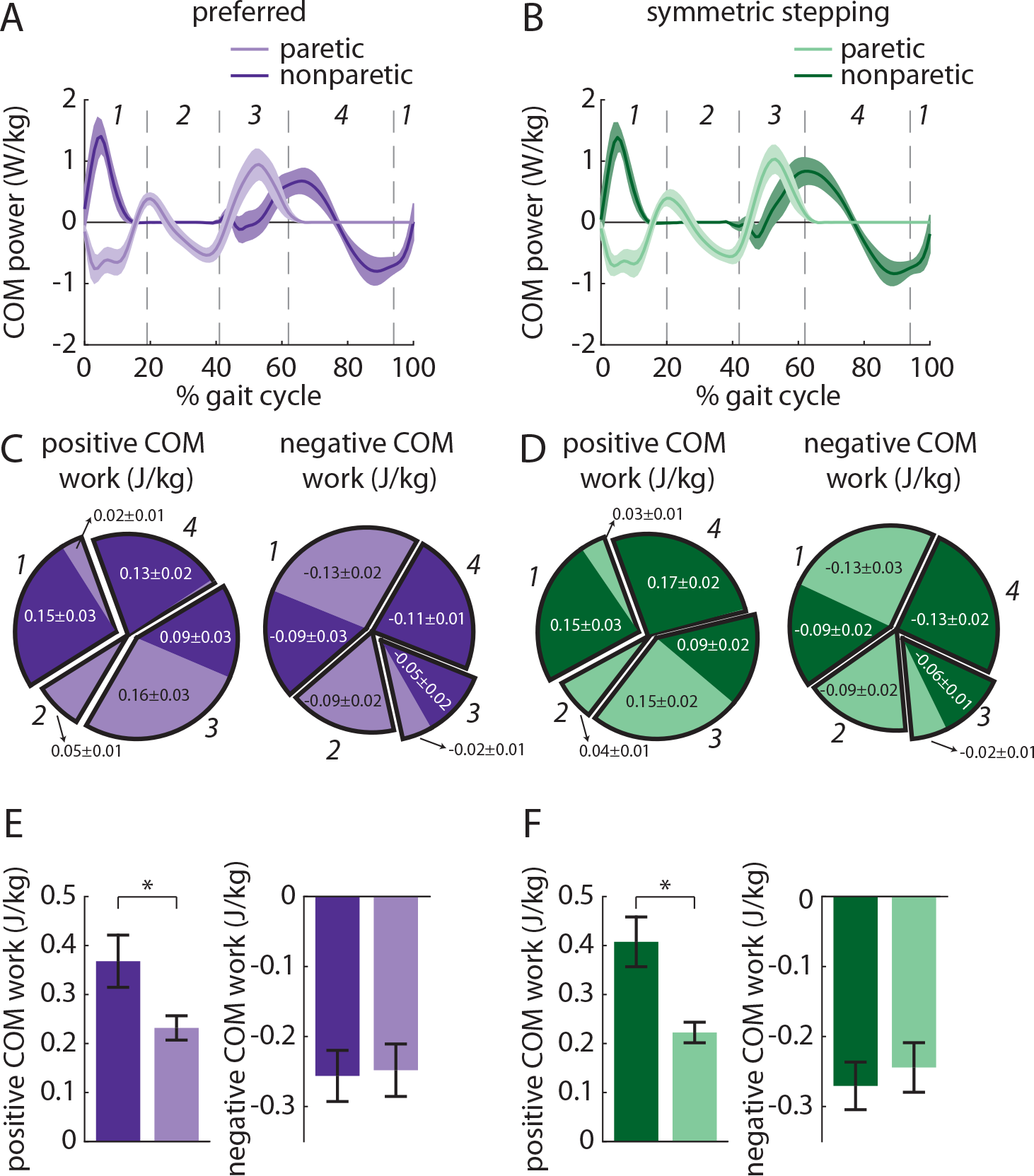
*A* and *B*) Center of mass (COM) power (mean ± SEM) generated by the paretic (light colors) and nonparetic (dark colors) limbs during *A*) preferred walking and *B*) symmetric stepping. The gait cycle is aligned to paretic heel-strike and partitioned into four periods defined by changes in direction of the COM velocity vector within the sagittal plane. *C* and *D*) Positive and negative COM work performed (mean ± SEM) by each limb in the four periods during *C*) preferred walking and *D*) symmetric stepping. Each pie chart displays fractions of overall positive or negative work during each period in each condition. Persons post-stroke perform more positive work during the fourth period (corresponding approximately with the transition from double support to nonparetic single support) during symmetric stepping as compared to preferred walking. Numerical labels on individual limb work contributing to less than 1% of total mechanical work in each condition and results of statistical analyses are omitted for clarity. *E* and *F*) Total positive and negative COM work (mean ± SEM; * p<0.05 between limbs) performed by each limb across all four periods during *E*) preferred walking and *F*) symmetric stepping. Persons post-stroke perform more positive work with the nonparetic limb than the paretic limb during both conditions.

ANOVA also revealed a significant main effect of leg on negative work done 1) by the paretic leg during the first period (step-to-step transition, nonparetic leg trailing) vs. the nonparetic leg during the third period (step-to-step transition, paretic leg trailing; F(1,9)=8.67, p=0.02), and 2) by the paretic leg during the third period vs. the nonparetic leg during the first period (F(1,9)=6.63, p=0.03). Post hoc analyses revealed that the paretic leg did significantly more negative work during the first period than the nonparetic leg did during the third period (p=0.02). However, the nonparetic leg did significantly more negative work during the first period than the paretic leg did during the third period (p=0.03).

We did not observe a significant main effect of condition on work done over any of the time periods (all p>0.07). We did observe a significant leg × condition interaction for the positive work done during the fourth period (nonparetic single support; F(1,9)=9.54, p=0.01). Post hoc analyses revealed that the nonparetic leg did significantly more positive work during the fourth period when symmetric stepping as compared to preferred walking (p=0.03).

A separate ANOVA revealed a significant main effect of leg on positive (but not negative; F(1,9)=0.15, p=0.71) work done across all time periods (F(1,9)=7.95, p=0.02). We did not observe a significant main effect of condition on positive or negative work done across all time periods (both p>0.30) but did observe a significant leg × condition interaction on positive work done across all periods (F(1,9)=5.26, p=0.048; F(1,9)=0.68, p=0.43 for negative work). Post hoc analyses revealed that the nonparetic leg did significantly more total positive work than the paretic leg (p=0.02).

### Less positive work done by the paretic leg is associated with higher cost of transport, slower walking, and increased interlimb asymmetry

We then assessed whether the positive (Figures 6A–6F) and negative work done by each leg across the gait cycle were related to cost of transport, gait speed, or interlimb asymmetry during preferred walking and symmetric stepping. Positive paretic work was significantly associated with decreased cost of transport (preferred walking: r=−0.84, p<0.01; symmetric stepping: r=−0.82, p<0.01; Figure 6A); positive nonparetic work was not significantly associated with cost of transport during either condition (both p>0.60, Figure 6B). Positive paretic work was also significantly associated with increased walking speed (preferred walking: r=0.90, p<0.01; symmetric stepping: r=0.84, p<0.01; Figure 6C) whereas positive nonparetic work was not (both p>0.63, Figure 6D). Finally, positive paretic work was significantly associated with decreased interlimb asymmetry during symmetric stepping (preferred walking: r=−0.60, p=0.06; symmetric stepping: r=−0.67, p=0.03; Figure 6E). Positive nonparetic work was not significantly associated with interlimb asymmetry during either condition (both p>0.48; Figure 6F). We did not observe significant associations between negative paretic or nonparetic work and cost of transport, walking speed, or interlimb asymmetry during either condition (all p>0.09).

**Figure 6.**
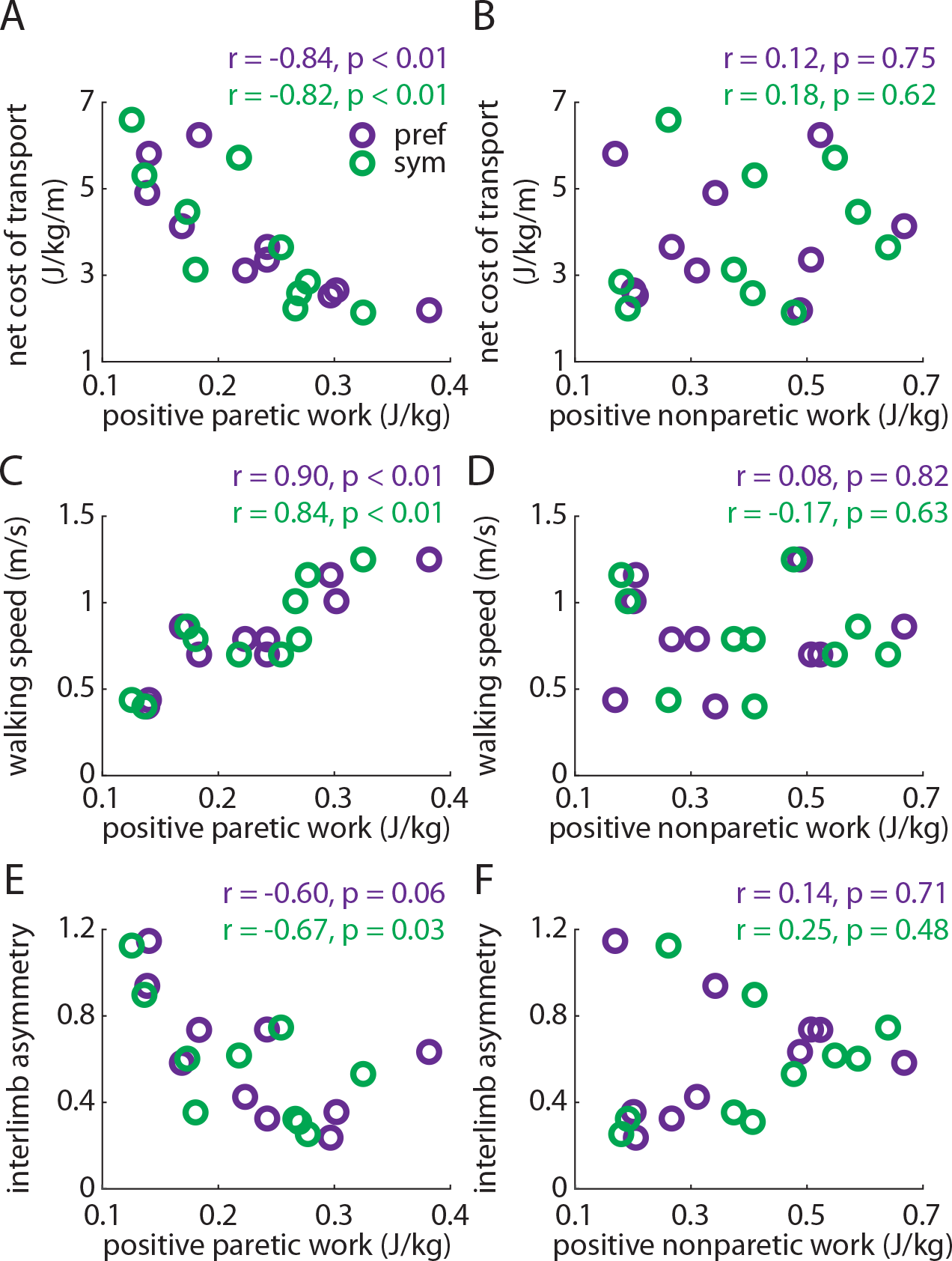
The amount of positive work performed by the paretic limb correlates strongly with the net metabolic cost of transport (*A*), walking speed (*C*), and interlimb asymmetry (*E*) during both preferred walking (purple) and symmetric stepping (green). The amount of positive work performed by the nonparetic limb is not significantly associated with any of these variables (*B*, *D*, and *F*) during either condition.

## DISCUSSION

In this study, we found that persons post-stroke restored step length symmetry using energetically expensive, asymmetric walking patterns. Persons post-stroke commonly exhibit asymmetric kinematics, impaired paretic propulsion during late paretic stance, and excessive compensatory vertical movement of the COM during nonparetic single support^4,31,32^. We found that these aberrant features persisted regardless of whether participants walked with symmetric or asymmetric step lengths. Deficits in positive paretic work were also unaffected by improvement in step length symmetry and were strongly associated with cost of transport, walking speed, and interlimb asymmetry. Our findings reveal that step length symmetry restoration does not necessarily result in positive changes elsewhere in the gait pattern after stroke. It is critical that interventions aiming to restore step length symmetry ensure that underlying impairments in gait mechanics are also addressed.

We do not intend for these findings to be interpreted as an indictment on step length asymmetry as a target of post-stroke gait rehabilitation. Interventions that target step length asymmetry have shown promise for improving gait speed^15^ and decreasing cost of transport^19^. However, step length asymmetry arises from a complicated series of deficits^4,5,22^. Our results emphasize that understanding the mechanisms underlying improvements in step length symmetry is critical to the success of these interventions. We propose that it is not so important *that* step length symmetry is restored, but rather *how* step length symmetry is restored that will facilitate gait improvement more broadly.

How should we design interventions to restore step length symmetry and decrease cost of transport? Improvement in the ability to generate positive paretic work is likely to result in a more symmetric, less effortful walking pattern. A substantial portion of the cost of transport can be attributed to the mechanical work generated by the legs on the COM during step-to-step transitions^26,33–35^. Persons post-stroke generate more positive work with the nonparetic leg than the paretic leg^4,27,28,31,36^, and this deficit is influenced by an inability to generate sufficient paretic propulsion during the step-to-step transition occurring during late paretic stance^32,37^. Interventions that improve paretic propulsion may then enhance the ability of the paretic leg to generate positive work and facilitate a faster, more symmetric, less costly gait pattern.

Why does the inability to generate positive paretic work increase the metabolic cost of walking after stroke? Prior studies have explained how the reduced positive work generated via paretic propulsion reverberates throughout the walking pattern. Paretic stance time is shortened relative to nonparetic stance^4^, and decreased paretic propulsion decreases the energy transmitted to the leg to initiate swing^31,38–40^. This necessitates compensatory mechanics to facilitate paretic leg swing. Often, vaulting (vertical movement of the COM generated primarily by the nonparetic leg) occurs to lift the paretic foot from the floor. This results in an elevated vertical COM velocity and increased positive work done to raise the COM (rather than direct the COM forward) during late paretic stance^8,27,28,31^. Persons post-stroke then often use a sequence of pelvic rotation, hip hiking, and hip circumduction to swing the leg and clear the foot^4,31^. This slow paretic leg swing extends nonparetic stance, increasing the positive nonparetic work done during single support^8^.

The inability to generate sufficient positive paretic work during late stance thus results in elevated metabolic cost of transport by 1) increasing positive nonparetic work during single support and late paretic stance, often in the vertical direction, and/or 2) slowing gait speed^41^ so that compensatory demands on the nonparetic leg are lessened. These altered mechanics persisted across both preferred walking and symmetric stepping, with the lone difference being that persons post-stroke generated more positive nonparetic work during single support when symmetric stepping (driven increased vertical COM velocity) to provide additional compensation during paretic swing. Fortunately, many interventions – including fast walking^42–44^, functional electrical stimulation of the plantarflexors^44^, and split-belt treadmill walking^20,45^ – show promise for improving paretic propulsion^37^. These interventions target paretic propulsion through combinations of improving ankle power generation and increasing paretic limb extension during late stance (i.e., trailing limb angle^39,46^). Improving propulsion by increasing ankle power is also a common goal in robotic designs^47–50^. Indeed, several of these interventions have resulted in improved step length symmetry^11,19^.

This study focused on the relationship between step length symmetry and cost of transport during only a single session of walking and only by using visual feedback to facilitate step length symmetry. Improved step length symmetry without changes in the underlying gait mechanics may affect cost of transport differently if the improved symmetry is achieved via long-term training. The participants in this study exhibited mild-to-moderate gait dysfunction and were in the chronic phase post-stroke. These results may not extrapolate to more impaired patients or patients in the acute phase post-stroke.

## CONCLUSIONS

Persons post-stroke restore step length symmetry using energetically expensive, asymmetric walking patterns. The similarity in cost of transport observed during preferred walking and symmetric stepping can be explained by the persistence of aberrant gait mechanics (specifically, impaired ability to generate positive paretic work) regardless of step length symmetry. Interventions that aim to restore step length symmetry are most likely to succeed when the gait mechanics that underlie the asymmetry are considered in the intervention design.

## ACKNOWLEDGMENTS

This project was funded by NIH grant R21 AG059184 to RTR and an AAP RREMS fellowship to SG. We thank Max Donelan for helpful suggestions on this project.

